# Single Cell-Type Spatial Proteomics Uncovers Regional Heterogeneity of Astrocytes

**DOI:** 10.64898/2026.03.30.715144

**Authors:** Chien-Chang Huang, Chiung-Yun Chang, Po-Chao Chan, Weng Man Chong, Hsiao-Jen Chang, Jung-Chi Liao

**Author notes:** Corresponding Author: Jung-Chi Liao - SYNCELL Inc., Taipei, Taiwan.

## Abstract

Astrocytes are a subset of glial cells in the central nervous system (CNS) that support numerous processes essential for brain function. Their functional diversity is thought to arise from specialized subpopulations with distinct molecular profiles. Although single-cell and single-nucleus RNA sequencing (scRNA-seq and snRNA-seq) have greatly advanced our understanding of astrocyte transcriptomic heterogeneity, mRNA abundance does not always correlate with protein levels because of post-transcriptional and translational regulation. Therefore, studying protein profiles remains essential to accurately capture astrocyte functional states and heterogeneity. Here, we used Microscoop Mint, a microscopy-guided spatial proteomics platform that integrates subcellular, region-specific sample preparation with LC-MS/MS-based mass spectrometry, enabling direct protein profiling of astrocytes in paraformaldehyde-fixed, optimal cutting temperature (OCT)-embedded mouse brain tissue. By applying this approach, we uncovered distinct regional-associated astrocyte proteomic signatures in the cerebral cortex and hippocampus and selected novel candidate protein markers for subsequent validation by immunofluorescence. Notably, MINK1 and PLEKHB1 showed preferential expression in hippocampal and cortical astrocytes, respectively, highlighting their potential as region-specific astrocyte markers. Overall, this strategy enables high-precision, unbiased spatial proteomic discovery at subcellular resolution, providing a powerful framework for linking molecular diversity to functional specialization in astrocyte biology.

## INTRODUCTION

Astrocytes are a subtype of glial cells and the most abundant cell type in the central nervous system (CNS)^1^. Far from being passive “support cells,” astrocytes are recognized as active participants in nearly every aspect of brain function and pathology. They carry out a wide array of essential functions including maintaining blood-brain barrier^2^, regulating brain homeostasis such as the supply of energy metabolites and neurotransmitter recycling^3^, regulating synapse formation, function, and elimination^4,5^, gliotransmission^6^, and contributing to CNS development and plasticity^7,8^. Although the brain contains numerous functionally distinct neuronal subtypes, astrocytes have been long considered a largely homogenous population^3,9,10^. In contrast to neurons, which can be classified by morphology, connectivity, or anatomical position, astrocytes appear relatively uniform across the CNS, making region-specific distinctions challenging to resolve^3^. Their lack of electrical excitability further obscured potential diversity, since astrocyte activity is not reflected by sodium channel-driven action potentials^11^. Recent advances in transcriptomics, new experimental tools, and growing recognition of astrocytes’ roles in brain circuits have highlighted their heterogeneity, yet the precise functions of distinct astrocyte subpopulations remain difficult to define^12^. Classical criteria for identifying cellular subtypes, such as anatomical features, molecular markers, developmental origins, and physiological properties, have been effective for categorizing neurons but are challenging to apply to astrocytes. Single-cell RNA-sequencing (scRNA-seq) and spatial transcriptomics methods have begun to address this gap by revealing astrocyte molecular diversity and identifying putative subpopulations across multiple brain regions, including the cortex and hippocampus^13^, cerebral cortex^14^, diencephalon^15^, frontal cortex and striatum^16^, and cortical gray matter as well as corpus callosum and cerebellar white matter^17^. Nonetheless, caution is warranted when inferring function solely from transcriptomic. A recent study has shown that key protein-level changes may not be captured by RNA expression data^18^, with large discrepancies between RNA and protein abundance reported in multiple tissues including the stomach, cerebral cortex, and cerebellum. Additionally, transcriptomically similar neurons can exhibit diverse morphologies and functions^19^, and the association between RNA expression and protein levels in astrocytes was previously reported to be relatively weak^20^. Thus, while scRNA-seq has been transformative for mapping astrocyte heterogeneity, it fails to capture functional states. Integrating transcriptomic data with protein-level analyses will be essential for accurately defining functionally distinct astrocyte subpopulations in the brain^14^. Spatial proteomic strategies are needed to directly define region- and niche-specific astrocyte protein signatures, motivating the development of methods capable of unbiased, high-resolution proteomic profiling within intact brain tissue. However, current platforms are frequently constrained by a fundamental trade-off between throughput, spatial resolution, multiplexing capacity, and analytical depth. Mass spectrometry-based imaging and multiplexed fluorescence imaging facilitate the mapping of dozens of targets at single-cell resolution^21,22,23^, while spatial barcoding technologies offer transcriptomic breadth that can be combined with targeted proteomics analysis. Yet, these remain ‘closed-system’ methodologies; they are inherently targeted, relying on validated antibodies or metal-tagged probes, and they frequently lack the subcellular precision required to isolate rare protein populations or discrete organelles. Such dependency precludes the discovery of novel biomarkers absent from pre-defined panels^24^.

To fill this gap, we previously developed the Microscoop technology^25,26^, a high-resolution, microscopy-guided spatial proteomics platform that integrates imaging-directed two-photon photo-biotinylation with downstream mass spectrometry analysis. This workflow enables selective labeling of proteins within user-defined regions across thousands of fields of view in an automated manner, allowing spatially resolved proteomic profiling directly from complex tissues. Here, we applied this platform to investigate regional astrocyte proteomic diversity in the cerebral cortex and hippocampus, where it enabled unbiased identification of distinct astrocyte proteomic signatures and the discovery of proteins enriched in specific anatomical locations.

## METHODS

### Animals

Brain tissue blocks from adult C57BL/6J mice (12 weeks old) were purchased from the Taiwan Mouse Clinic (TMC). All animal procedures were conducted in accordance with protocols approved by the Institutional Animal Care and Use Committee (IACUC) of the Biomedical Translation Research Center, Academia Sinica, Taiwan. Briefly, mice were anesthetized and transcardially perfused with 0.9% saline to remove circulating blood, followed by 4% paraformaldehyde (PFA, P6148, Sigma-Aldrich, MO, USA) for tissue fixation. Brains were carefully dissected, embedded in optimal cutting temperature (OCT) compound (3801480, Leica Biosystems Nussloch GmbH, Nubloch, Germany), rapidly frozen and stored at -80 freezer. Frozen brain blocks were sectioned at a thickness of 10 μm using a cryostat for subsequent analysis.

### Immunostaining of Astrocyte and Novel Candidate Proteins Validation

Frozen mouse brain sections were permeabilized and blocked overnight at 4 in PBS containing 1% goat serum (16210072, Thermo Fisher Scientific, MA, USA) and 0.5% Triton X-100 (T8787, Sigma-Aldrich, MO, USA). Sections were subsequently incubated with primary antibodies including mouse anti-GFAP (clone GA5, MAB360, Sigma-Aldrich, MA, USA) for Microscoop photolabeling, or other antibodies for novel candidate proteins validation (Table S1), diluted 1:500 in PBS containing 0.1% Triton X-100 (PBST), overnight at 4°C. After multiple washes by PBST, samples were incubated overnight at 4°C with Alexa Fluor 568-conjugated donkey anti-mouse secondary antibody or Alexa Fluor Plus 647-conjugated donkey anti-rabbit secondary antibody (A10037 and A32795TR, Thermo Fisher Scientific, MA, USA), diluted 1:500 in PBS. Following PBST washes, stained tissue sections were stored in PBS at 4°C until photolabeling or imaged with confocal microscope (LSM 880, Carl Zeiss AG, Oberkochen, Germany).

### Microscoop Photolabeling

For each replicate, 18 mouse brain coronal sections were stained with anti-GFAP antibody (a canonical astrocyte marker). Six sections were used for photolabeling of astrocytes in cortex and that in hippocampus respectively, the remaining six sections were kept in dark as unlabeled controls. All samples were processed according to the Synlight-Rich Kit protocol (SYN-RI0106, Syncell, Taipei, Taiwan). Briefly, the GFAP-staining tissue slices were sequentially blocked using Block1 and Block2 blocking reagents. To avoid evaporation of photolabeling reagent, leak-proof wells are created around each sample using imaging spacers (iSpacer IS201/IS211, SunJin Lab, Hsinchu, Taiwan). The sections were then incubated in photoreactive probes and overlayed with cover glass. Image acquisition was performed using the Microscoop MINT system (model MS.1011, Syncell, Taipei, Taiwan). Regions of interest (ROIs) within each field of view (FOV) were identified and converted into binary masks using Autoscoop software (version 1.0.1.11, Syncell, Taipei, Taiwan). Binarization thresholding and morphological operation with erosion were applied to refine the high intensity regions, after which the masks guided two-photon laser illumination of astrocyte-specific ROIs, triggering photo-biotinylation through a light-sensitive probe activation. This automated photolabeling procedure was repeated across hundreds to thousands of FOVs per sample. Following photolabeling, the photoreactive probes were removed, and the brain sections were thoroughly washed with PBST to minimize interference from unreacted probes during subsequent pulldown assays. Verification of biotinylation following photolabeling was performed. Sections were incubated with the verification buffer from the Synlight-Rich Kit (SYN-RI0106, Syncell, Taipei, Taiwan) for three hours at room temperature. After three PBST washes, nuclei were counterstained by DAPI (D1306, Thermo Fisher Scientific, MA, USA). Imaging was performed using Microscoop with the DAPI/FITC/TRTIC/Cy5 channels for epifluorescence imaging.

### Liquid Chromatography-Tandem Mass Spectrometry (LC-MS/MS) Analysis

Biotinylated proteins were purified using Synpull Kit (SYN-PU0106, Syncell, Taipei, Taiwan) according to the manufacturer’s instructions and subsequently analyzed to mass spectrometry, as detailed in Lin (2025)^26^. Purified photolabeled/un-labeled peptides were analyzed on an Orbitrap Fusion Lumos Tribrid quadrupole-ion trap-Orbitrap mass spectrometer (Thermo Fisher Scientific, MA, USA) equipped with UltiMate 3000 RSLCnano system (Thermo Fisher Scientific, MA, USA). Desalted and dried peptides were reconstituted in 0.1% formic acid (FA) in water and separated on a 75 µm × 250 mm column (pore size 130 Å, particle size 1.7 µm, nanoEase M/Z Peptide CSH C18, Waters). The mobile phase A consisted of 0.1% FA in water, and mobile phase B was 100% acetonitrile (ACN) with 0.1% FA. Peptides were separated using a linear gradient from 2% to 38% ACN with 0.1% FA over 120 min at a flow rate of 300 nL/min. The mass spectrometer was operated in data-independent acquisition (DIA) mode. Full MS spectra were acquired in the Orbitrap at a resolution of 120,000 with a scan range of 375–1000 m/z, an automatic gain control (AGC) target of 1×10^6^, and a maximum injection time of 50 ms. For DIA MS/MS, 40 scan events were performed with a 10 m/z isolation window, fragmentating ions within the m/z range of 400–800. MS/MS scans were acquired in high-energy collision dissociation (HCD) mode at 30% normalized collision energy, with an AGC target of 4×10^5^ and a maximum injection time of 54 ms. Fragment ions were scanned in Orbitrap with a resolution of 30,000.

### Mass Spectrometry Data Analysis

Raw DIA files from both photolabeled and unlabeled (control) samples were analyzed simultaneously using Spectronaut (version 19.6.250122.62635; Biognosys AG, Schlieren, Switzerland) in directDIA™ mode, which enables spectral library-free analysis. Protein identification was performed using the Pulsar search engine against the UniProtKB/Swiss-Prot mouse proteome database (version 2023_04). Search parameters were defined as follows: variable modifications included methionine oxidation and N-terminal protein acetylation, while carbamidomethylation of cysteine residues was set as a fixed modification. Trypsin/P was specified as the digestion enzyme, allowing up to two missed cleavages. Label-free quantification was based on MS1-level peptide peak intensities, and protein abundance was estimated by averaging the top three most abundant peptides per protein. Missing values were imputed using a global imputation strategy. For statistical analysis, unpaired t-tests were conducted to assess differential protein abundance between photolabeled and unlabeled controls. Volcano plots were generated to visualize fold changes against significance p-value. Proteins with a fold change ≥ 1.5 and p ≤ 0.05 were considered candidates associated with the regions of interest (ROIs). The astrocyte database was obtained from UniProt and the Human Protein Atlas using *Mus musculus* as the target species. Known astrocyte proteins were identified by searching and cross-referencing a combination of the keywords including “astrocyte,” “hippocampus,” and “cortex.” From the cross-referenced database, 352 astrocyte marker proteins were identified. The full list is available as supplementary table S4.

### Selection of Region-Specific Protein Candidates

Venn diagram analysis was used to identify proteins consistently identified in two independent biological replicates, establishing each foundational proteomic datasets for cortical and hippocampal astrocytes. These datasets were further refined using a stringent sequential filtering criteria: (1) a fold-change (FC) ≥ 1.5 in protein abundance relative to the unlabeled (UL) control group, (2) a statistically significant p-value ≤ 0.05, and (3) a minimum of two unique peptides per protein identification. Subsequent comparative Venn analysis was performed on these filtered datasets to delineate region-specific proteomes for the cortex and hippocampus.

## RESULTS AND DISCUSSION

We applied the Microscoop, an imaging-guided proteomic sample preparation system that integrates microscopy and photochemistry, to biotinylate proteins within the defined regions of interest (ROIs). A workflow coupling Microscoop with mass spectrometry (MS) enables unbiased proteomic profiling at single-cell or subcellular resolution^25,26^. As illustrated in Figure 1a, the Microscoop-to-MS workflow consists of five steps: (1) sample preparation, (2) patterned photo-biotinylation, (3) protein extraction, (4) enrichment and digestion, and (5) proteomic identification and analysis. The target ROI in fixed cells or tissue sections is visualized by a fluorescent signal, and a photo-reactive biotin reagent is added to the sample. Using built-in image-processing algorithms, binary masks are generated to define the ROIs and to guide illumination of a femtosecond two-photon laser on the sample across hundreds to thousands of fields of view automatically. Once illuminated, the photo-reactive biotin inside the sample is activated at pixel-level precision, enabling spatially restricted protein biotinylation. Samples are finally collected and lysed; biotinylated proteins are enriched via streptavidin beads and analyzed by LC-MS/MS to identify localization-specific proteomes. In this study, astrocytes in mouse brain cryo-sections were immunostained against glial fibrillary acidic protein (GFAP), a canonical astrocytic marker^27^. Astrocytes in the cortex and hippocampus were identified, segmented, and selectively biotinylated using the Microscoop platform (Figure 1a). Biotinylation precision was confirmed by the colocalization of biotinylation signals within astrocytes in both regions (Figure 1b). Following Microscoop labeling, biotinylated astrocytes from photolabeled mouse brain tissues were purified and analyzed by LC-MS/MS (Figure 1a). The same set of samples without being exposed to light were also prepared in parallel as unlabeled control. Using this approach, we identified the proteome of astrocytes in the cerebral cortex (Figure 2a) and hippocampus (Figure 2b). Mass data from two independent biological replicates were filtered using a stringent sequential thresholding of fold change (photolabeled / unlabeled control) ≥ 1.5, p value ≤ 0.05 and unique peptide ≥ 2. This analysis identified a total of 1,803 proteins significantly enriched in the photolabeled astrocytes compared to the unlabeled controls in the cerebral cortex (Table S2), and 987 enriched proteins in the hippocampus (Table S3). The protein datasets were cross-referenced against a curated database comprising 352 canonical astrocyte markers (Table S4). Seventy-one and forty-seven pan-astrocyte markers were found in the cortex (Figure 2a) and hippocampus (Figure 2b) datasets respectively, with 43 shared between both regions. These shared proteins encompassed several key functional molecules including glutamate transporter 1 (SLC1A2) and glutamate–aspartate transporter (SLC1A3) for neurotransmitter uptake and recycling; the potassium channel (KCNJ10), aquaporin-4 (AQP4), and sodium/potassium-transporting ATPase subunit beta-2 (ATP1B2) for ion homeostasis and water balance; GFAP, vimentin, radixin (RDX) and microtubule-associated protein tau (MAPT) for structural integrity; gap junction alpha-1 protein (GJA1) for intercellular communication; aldehyde dehydrogenase 1 family member L1 (ALDH1L1) for folate metabolism; apolipoprotein E (APOE), fatty acid-binding protein 7 (FABP7) and glycerol kinase (GK) for lipid metabolism; and lactate dehydrogenase A (LDHA) for glucose metabolism. Collectively, these proteins represent core components of astrocytic physiology and are widely recognized as fundamental to astrocyte biology^28,29^. These results demonstrate the Microscoop platform’s efficacy in capturing a broad spectrum of established markers that define the core physiological and functional identity of astrocytes across both the cortex and hippocampus.

**Figure 1.**
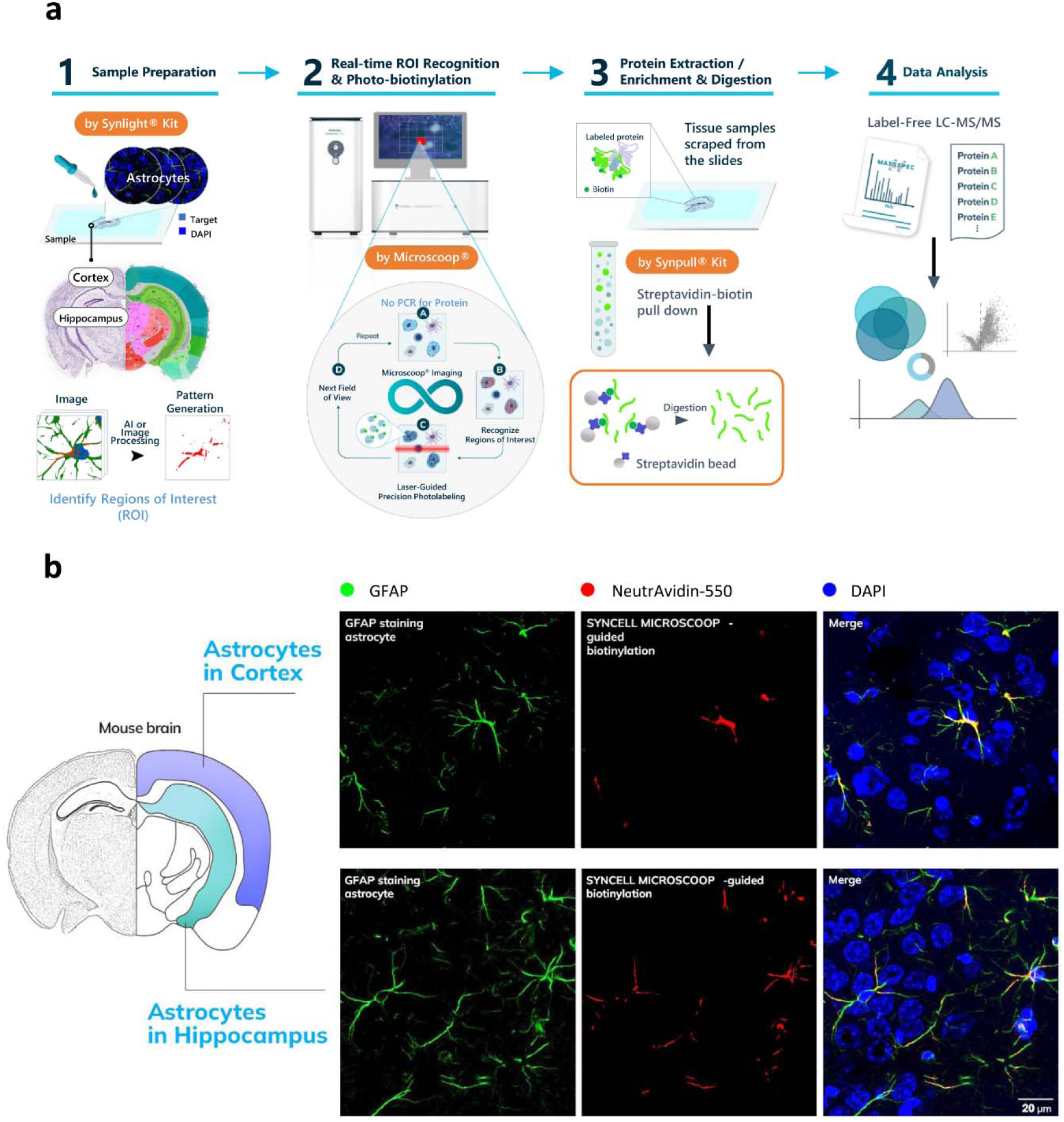
Image-guided biotinylation and proteomic identification of cortical and hippocampal astrocytes in mouse brains. (a) Microscoop Mint workflow consists of five steps: (1) Sample preparation: mouse brain tissues were stained with the astrocytic marker GFAP to delineate ROIs, which were annotated and converted into binary image masks by Autoscoop software. (2) Patterned photo-biotinylation: femtosecond two-photon illumination triggered spatially confined biotinylation across hundreds to thousands of FOVs. (3) Protein extraction: samples from multiple slides were pooled to increase protein yield. (4) Enrichment and digestion: biotinylated proteins were selectively isolated using streptavidin beads and digested into peptides. (5) Proteomic identification: peptides from labeled and unlabeled samples were analyzed by LC-MS/MS, yielding location-specific proteomes with high resolution and sensitivity. (b) Epifluorescence imaging of cortical and hippocampal astrocytes and validation of Microscoop-guided biotinylation. Cortical and hippocampal astrocytes in coronal sections of C57BL/6J mouse brain were stained using anti-GFAP antibody, and were then recognized, and photolabeling by Microscoop platform. Biotinylated proteins were validated by DyLight 550-conjugated neutravidin protein (NeutrAvidin-550) and cell nuclei were stained by DAPI. Scale bar, 20 μm.

**Figure 2.**
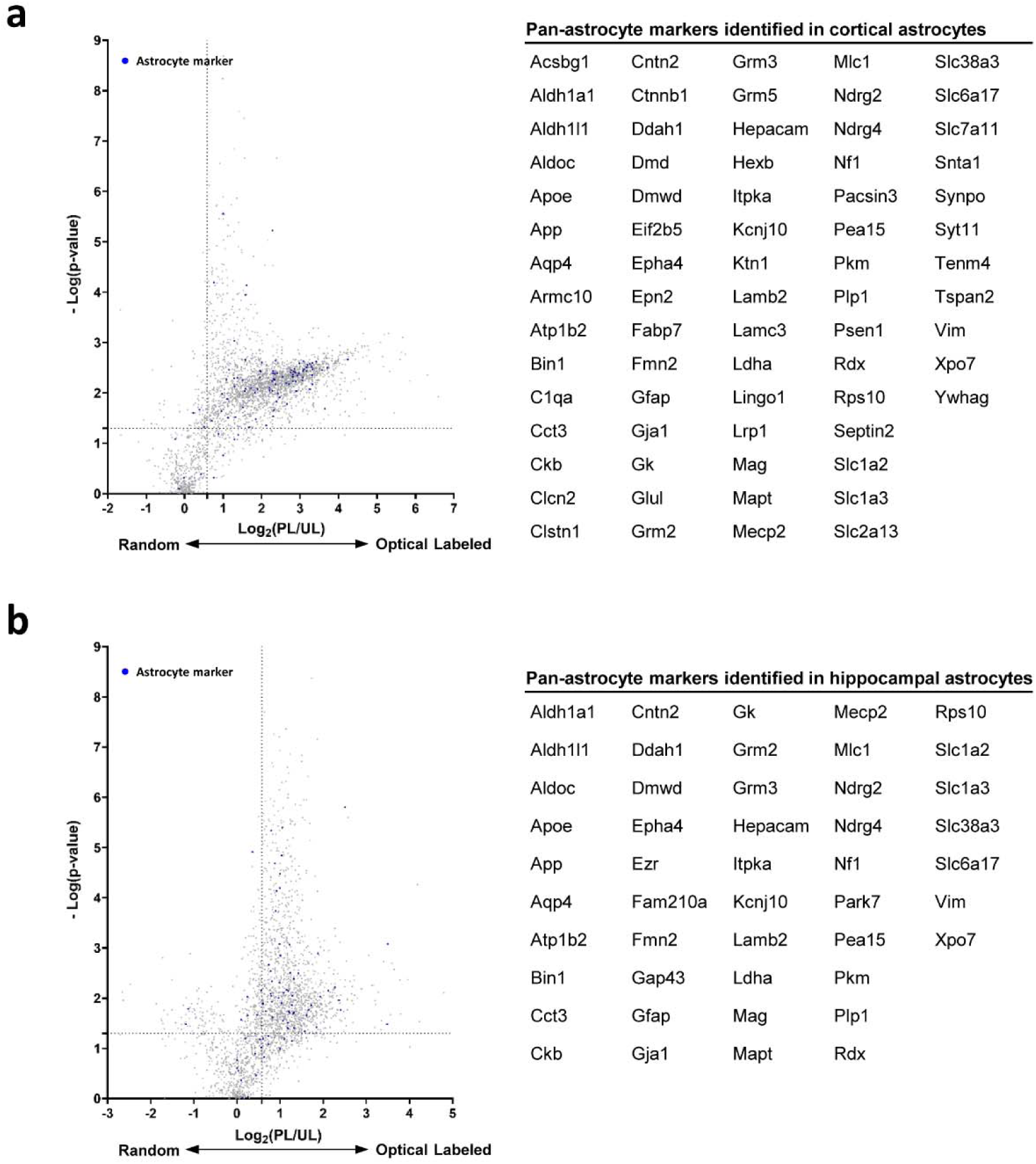
Proteome identification of cortical and hippocampal astrocytes. Volcano plots illustrate the differential enrichment of biotinylated proteins captured from cortical (a) and hippocampal (b) astrocytes. The x-axis denotes the log_2_ fold-change (FC) in protein abundance, comparing the photolabeled (PL) group to the unlabeled (UL) control group. The y-axis represents statistical significance as -log_10_(p-value). Proteins annotated as canonical astrocyte markers are highlighted in blue. Candidate regional markers were prioritized using stringent filtering criteria: fold-change ≥ 1.5, p-value ≤ 0.05, and a minimum of ≥ 2 unique peptides per protein.

Having established the proteomic profiles of cortical and hippocampal astrocytes, we compared the proteomes between the two regions. To investigate the regional heterogeneity of the astrocyte proteome, we analyzed cortex and hippocampus proteomes, each comprising two independent biological replicates (BR) datasets (Figure 3a). Initial intra-regional analysis using Venn diagrams demonstrated high reproducibility between replicates, yielding 1,594 high-confidence protein IDs in the cortex (Table S5) and 1,228 in the hippocampus (Table S6). Comparative inter-regional analysis revealed a core astrocytic proteome consisting of 314 proteins shared between both areas. Beyond this common signature, we identified significant regional divergence, with 226 proteins uniquely expressed in the cortex (Table S7) and 128 proteins specific to the hippocampus (Table S8). In cortical astrocytes, we observed significant up-regulation of structural and extracellular matrix-related proteins, including cystatin-C (CST3), fibulin-5 (FBLN5), laminin subunit gamma-3 (LAMC3), and alongside the signaling adaptor pleckstrin homology domain-containing family B member 1 (PLEKHB1) (Figure 3b). In addition, cortical astrocytes showed significantly higher enrichment of cytosolic 10-formyltetrahydrofolate dehydrogenase (ALDH1L1), aldehyde dehydrogenase 1A1 (ALDH1A1), laminin subunit alpha-2 (LAMA2), laminin subunit alpha-5 (LAMA5), LAMC3, collagen alpha-1(I) chain (COL1A1), and collagen alpha-2(I) chain (COL1A2), highlighting their strong involvement in the astrocyte’s role as a metabolic hub and brain’s structural integrity (Table S7). Conversely, hippocampal astrocytes were characterized by the enrichment of coactosin-like protein (COTL1), misshapen-like kinase 1 (MINK1), and PH and SEC7 domain-containing protein 3 (PSD3) (Figure 3b). Moreover, hippocampal astrocytes showed elevated levels of neurotransmitter transporters and receptors including the γ-aminobutyric acid (GABA) transporter (SLC6A11), and sodium-dependent glutamate/aspartate transporter 2 (SLC1A2), consistent with their role in neurotransmitter homeostasis (Table S8).

**Figure 3.**
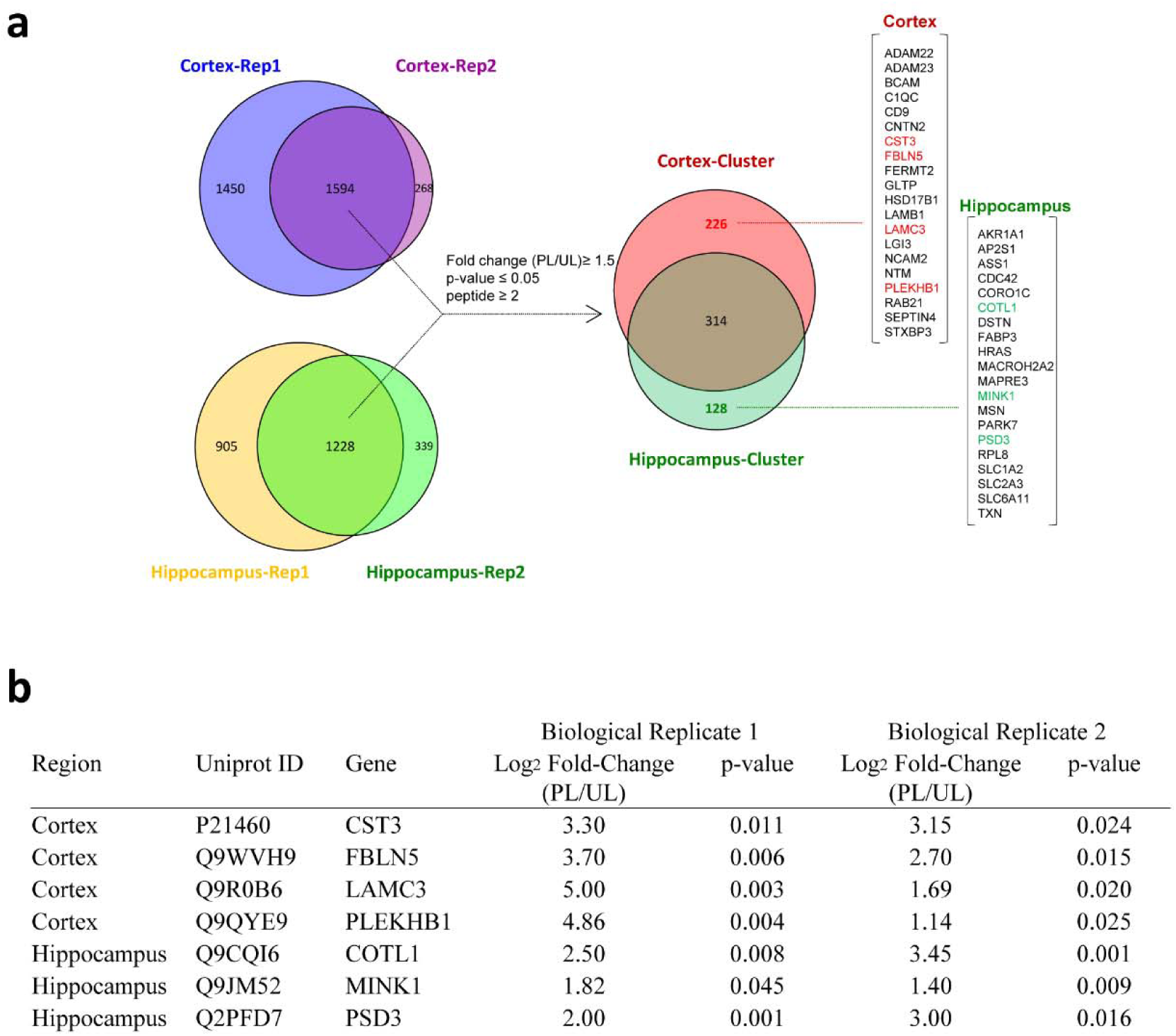
Region-enriched proteins in cortical and hippocampal astrocytes. (a) Venn diagram analysis depicting the overlap of protein datasets between two independent biological replicates (Rep1 and Rep2) for cortical and hippocampal regions. Following the application of stringent filtering criteria, a secondary comparative analysis was performed to isolate region-specific proteomes. Candidate proteins selectively enriched in the cortex (red) or hippocampus (green) were prioritized for subsequent validation. (b) Representative candidate proteins exhibiting significant differential expression in cortical or hippocampal astrocytes across two independent biological replicates. For each replicate, values represent the log_2_ (fold-change) in protein abundance, comparing the photolabeled (PL) group to the unlabeled (UL) control group. Statistical significance was determined via Student’s t-test, with p ≤ 0.05 considered significant.

To validate region-specific astrocyte candidate markers, we selected a subset of proteins that demonstrated significant and reproducible enrichment (log_2_ fold-change ≥ 1.1; p ≤ 0.05) in either the cortex or hippocampus across two independent biological replicates (Figure 3b). These candidates exhibited consistent enrichment profiles, underscoring their reliability as region-specific markers. To confirm their spatial distribution, we performed immunostaining and confocal imaging of the candidates together with GFAP in both the cortical and hippocampal regions of the brain tissue. Aligning with previous single-cell RNA sequencing (scRNA-seq) datasets^13^, GFAP was ubiquitously detected in astrocytes across both the cortical and hippocampal regions (Figures 2, 4, and S1) consistent with its established role as a canonical pan-astrocytic marker^27^ and as a representative astrocyte marker used in this study. In consistency with our proteomic findings, MINK1 and PLEKHB1 exhibited distinct regional enrichment. MINK1 was confined to mostly hippocampal astrocytes but not cortical astrocytes, while PLEKHB1 was found dominantly in cortical astrocytes but not hippocampal astrocytes (Figure 4a and 4b). Although neither protein has previously been linked to astrocyte biology, their localized expression suggests that they may serve as promising candidates for exploring region-specific astrocyte function within the central nervous system.

**Figure 4.**
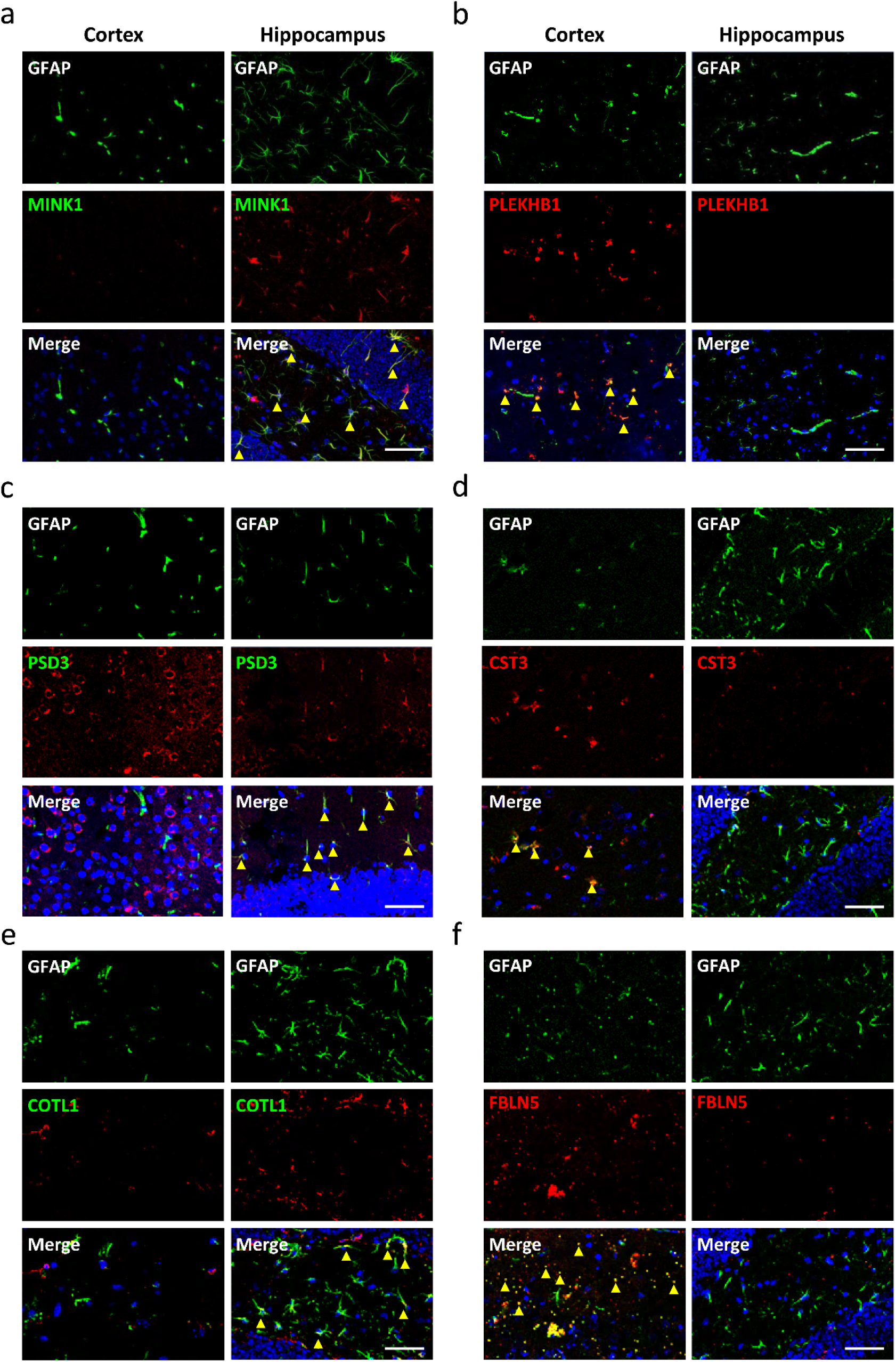
Validation of region-specific astrocyte candidates via confocal microscopy. Representative confocal images showing the expression and localization of selected protein candidates within cortical and hippocampal astrocytes. Astrocytes were identified by co-staining with the canonical marker GFAP (green) and DAPI for nuclei (blue). Hippocampus-specific candidates (green font) are (a) MINK1, (c) PSD3, and (e) COTL1. Cortex-specific candidates (red font) are (b) PLEKHB1, (d) CST3, and (f) FBLN5. Yellow arrowheads indicate the colocalization of GFAP with the respective protein markers. Scale bar, 50 μm.

Another notable finding is the distinct astrocyte expression of PSD3 and COTL1 in hippocampus (Figure 4c and 4e). Although previous in situ hybridization studies have documented broad PSD3 mRNA expression across the cerebral cortex, thalamus, and hippocampal pyramidal layers^30^, so as our immunostaining confirming ubiquitous PSD3 protein expression in both the cortex and hippocampus of the brain, our analysis revealed a sharp regional dichotomy in its cellular localization: PSD3 colocalized robustly with GFAP in the hippocampus but was excluded from cortical astrocytes (Figure 4c). In terms of molecular function, PSD3 acts as a guanine nucleotide exchange factor (GEF) for ADP-ribosylation factor 6 (ARF6), a small GTPase that orchestrates membrane trafficking and actin cytoskeletal remodeling^31^. Although the precise role of astrocytic PSD3 remains to be investigated, its enrichment may suggest that hippocampal astrocytes may be specialized for dynamic cytoskeletal remodeling to support synaptic plasticity and neurogenic niches.

In contrast, a distinct set of proteins, CST3 (Figure 4d), FBLN5 (Figure 4f), and LAMC3 (Figure S1), displayed colocalization with GFAP exclusively in the cortex. FBLN5 is an extracellular matrix glycoprotein essential for the assembly of elastic fibers^32^. Its cortical-exclusive expression suggests that cortical astrocytes may maintain a specialized, structurally resilient microenvironment. Furthermore, LAMC3 is one of components of laminins which are major components of the basal lamina^33^ and is also a critical component of the cortical pial basement membrane that serves as an essential structural scaffold for Cajal-Retzius and radial glial cell^34^. The enrichment of LAMC3 in cortical but not hippocampal astrocytes suggests these cells may actively sustain the unique pial-glial basement membrane or blood-brain barrier (BBB) architectures of the neocortex.

Collectively, our findings support an astrocyte functional dichotomy may exist across brain areas: while hippocampal astrocytes may be specialized for the high synaptic turnover associated with neurogenesis, cortical astrocytes may prioritize the maintenance of a robust ECM scaffold. Our findings highlight the profound molecular heterogeneity of astrocyte populations and establish a framework for investigating their regional specialization in brain physiology in future studies.

## CONCLUSION

Together, these results indicate that an imaging-guided proteomics approach can enable unbiased, high-resolution identification of common and region-specific astrocyte protein signatures in the cerebral cortex and hippocampus. The observed differences in protein expression of a subset of markers reveal anatomic astrocyte heterogeneity within complex brain tissue environments.

---

**Supplementary Figure S1.**
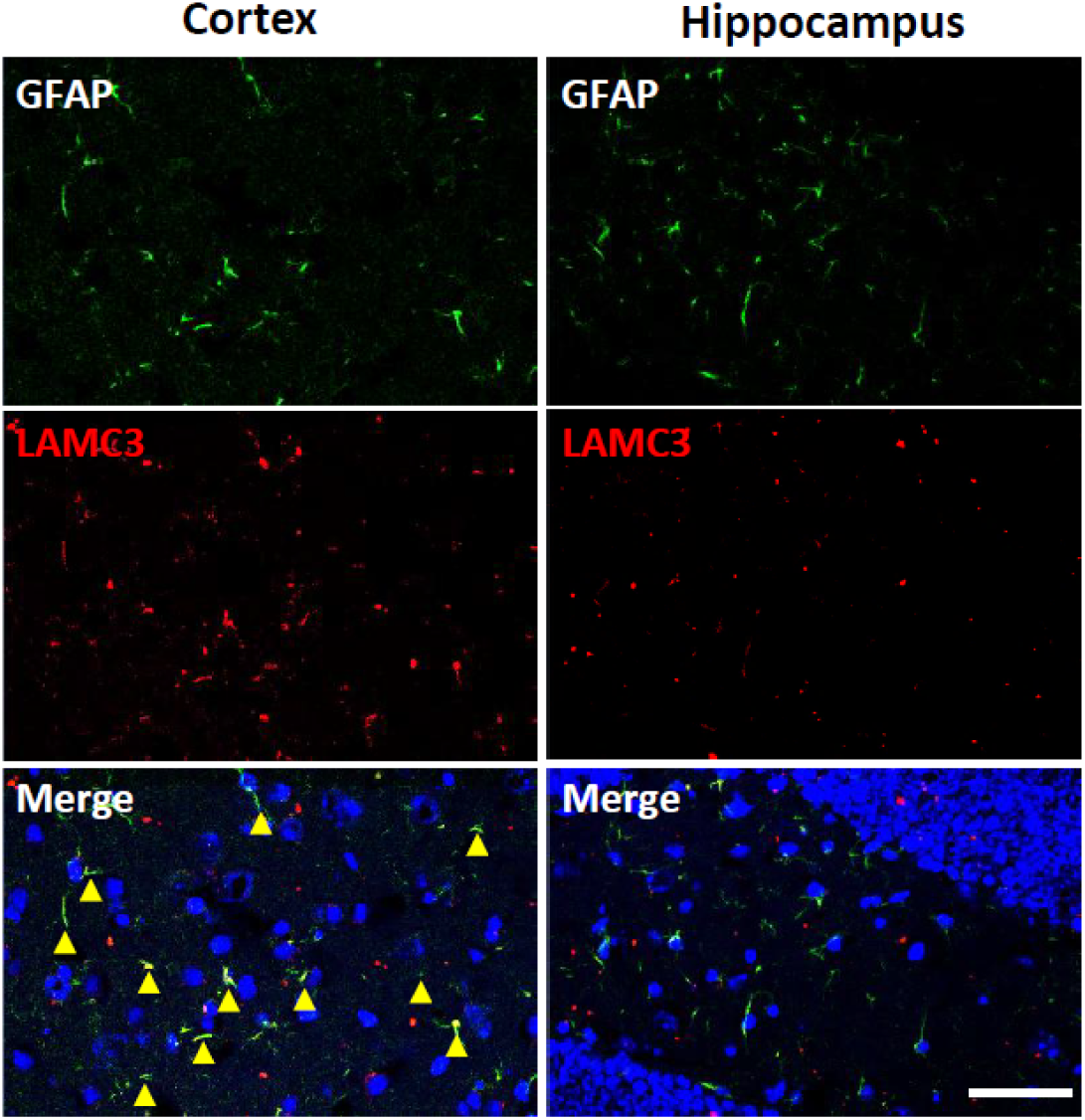
Validation of a cortical astrocyte candidate LAMC3 via confocal microscopy. Representative confocal images showed the expression and localization of LAMC3 within cortical and hippocampal astrocytes. Astrocytes were identified by co-staining with the canonical marker GFAP (green) and DAPI for nuclei (blue). Yellow arrowheads indicate the colocalization of GFAP with LAMC3. Scale bar, 50 μm.

## ASSOCIATED CONTENT

## Data Availability Statement

The mass spectrometry datasets have been deposited to the MassIVE (https://massive.ucsd.edu/ProteoSAFe/private-dataset.jsp?task=d34d25a6b7b2406fabee042be5f24f22) via the MassIVE partner repository with the data set identifier PXD072480 and 10.25345/C5XW48916.

## Supporting Information

Table S1: List of antibodies used for immunofluorescent staining (PDF).

Table S2: Comprehensive proteome of cortical astrocytes (XLSX).

Table S3: Comprehensive proteome of hippocampal astrocytes (XLSX).

Table S4: Curated database of canonical astrocyte marker proteins (XLSX).

Table S5: Commonly identified proteins across two cortical astrocyte replicates (XLSX).

Table S6: Commonly identified proteins across two hippocampal astrocyte replicates (XLSX).

Table S7: Proteins enriched specifically in cortical astrocytes (XLSX).

Table S8: Proteins enriched specifically in hippocampal astrocytes (XLSX).

## AUTHOR INFORMATION

## Authors

Chien-Chang Huang - SYNCELL Inc., Taipei, Taiwan;

Chiung-Yun Chang - SYNCELL Inc., Taipei, Taiwan;

Po-Chao Chan - SYNCELL Inc., Taipei, Taiwan;

Weng Man Chong - SYNCELL Inc., Taipei, Taiwan;

Hsiao-Jen Chang- SYNCELL Inc., Taipei, Taiwan;

## Author Contributions

Conceptualization: C.C.H., W.M.C.; methodology: C.C.H., H.J.C., W.M.C.; Investigation : C.C.H., H.J.C., Validation: C.C.H.; Data curation: C.C.H., H.J.C., C.Y.C.; Data analysis: : C.C.H, C.Y.C., W.M.C., P.C.C.; Visualization: C.C.H., C.Y.C., P.C.C., W.M.C.; writing- original draft preparation: P.C.C., : C.C.H, C.Y.C.; writing-review and editing: P.C.C., : C.C.H, C.Y.C., W.M.C.; project administration: W.M.C.; Supervision: J.C.L.; Resources: J.C.L.; and funding acquisition: J.C.L.. All authors have given approval to the final version of the manuscript.

## NOTES

Patent applications related to the subject matter of this publication have been filed. All authors declare that they are current employees of Syncell Inc.

## ACKNOWLEDGMENT

We thank the Core Facilities of Translational Medicine of BioTReC (National Biotechnology Research Park, Academic Sinica), Taiwan for the technical support and data analysis, and Professor Hsueh-Fen Juan at National Taiwan University for the useful discussion, advice, and proofread the manuscript.

## ABBREVIATIONS

ROI: region of interest
FOV: field of view
GFAP: glial fibrillary acidic protein
CNS: central nervous system
DIA: data-independent acquisition
PBS: phosphate buffered saline
PBST: phosphate buffered saline containing 0.1% Triton X-100
DAPI: 4′,6-diamidino-2-phenylindole.

