## Supplementary material for "Single Cell-Type Spatial Proteomics Uncovers Regional Heterogeneity of Astrocytes": Table S1: Table S1_List of antibodies used for immunofluorescent staining (PDF).pdf

Supplementary Table 1

| Antibodies | Source | Identifier |
|---|---|---|
| Rabbit polyclonal anti-COTL1 | Proteintech | Cat# 10781-1-AP,<br>RRID: AB_2084785 |
| Rabbit polyclonal anti-PSD3 | Novus | Cat# NBP2-55678,<br>RRID: AB_3340881 |
| Rabbit polyclonal anti-MINK1 | Proteintech | Cat# 13137-1-AP,<br>RRID: AB_10644151 |
| Mouse monoclonal anti-LAMC3 | Proteintech | Cat# 67261-1-Ig,<br>RRID: AB_2882532 |
| Rabbit polyclonal anti-Cystatin C | Proteintech | Cat# 12245-1-AP,<br>RRID: AB_2088058 |
| Rabbit polyclonal anti-PLEKHB1 | Proteintech | Cat# 10512-1-AP,<br>RRID: AB_2166943 |
| Rabbit polyclonal anti-FBLN5 | Proteintech | Cat# 12188-1-AP,<br>RRID: AB_2105939 |
| Rabbit polyclonal anti-GFAP | Proteintech | Cat# 81063-1-RR,<br>RRID: AB_2923699 |
| Mouse monoclonal anti-GFAP | Merck Millipore | Cat# MAB360,<br>RRID: AB_11212597 |
| Donkey anti-Rabbit IgG,<br>Alexa Fluor Plus 555 | Thermo Fisher Scientific | Cat# A32794,<br>RRID: AB_2762834 |
| Donkey Anti-Mouse IgG,<br>Alexa Fluor 568 | Thermo Fisher Scientific | Cat# A10037,<br>RRID: AB_11180865 |
| Donkey anti-Rabbit IgG,<br>Alexa Fluor Plus 647 | Thermo Fisher Scientific | Cat# A32795TR,<br>RRID: AB_2866496 |
| Donkey anti-mouse IgG,<br>Alexa Fluor 647 | Thermo Fisher Scientific | Cat# A-21235,<br>RRID: AB_2535804 |
